# Identification of novel bat coronaviruses sheds light on the evolutionary origins of SARS-CoV-2 and related viruses

**DOI:** 10.1101/2021.03.08.434390

**Authors:** Hong Zhou, Jingkai Ji, Xing Chen, Yuhai Bi, Juan Li, Tao Hu, Hao Song, Yanhua Chen, Mingxue Cui, Yanyan Zhang, Alice C. Hughes, Edward C. Holmes, Weifeng Shi

**Affiliations:** Key Laboratory of Etiology and Epidemiology of Emerging Infectious Diseases in Universities of Shandong, Shandong First Medical University & Shandong Academy of Medical Sciences, Taian 271000, China; Landscape Ecology Group, Center for Integrative Conservation, Xishuangbanna Tropical Botanical Garden, Chinese Academy of Sciences, Menglun, Mengla, Yunnan 666303, China; CAS Key Laboratory of Pathogenic Microbiology and Immunology, Institute of Microbiology, CAS Center for Influenza Research and Early-warning (CASCIRE), CAS-TWAS Center of Excellence for Emerging Infectious Diseases (CEEID), Chinese Academy of Sciences, Beijing 100101, China; Research Network of Immunity and Health (RNIH), Beijing Institutes of Life Science, Chinese Academy of Sciences, Beijing 100101, China; Center of Conservation Biology, Core Botanical Gardens, Chinese Academy of Sciences, Menglun, Mengla, Yunnan 666303, China; Marie Bashir Institute for Infectious Diseases and Biosecurity, School of Life and Environmental Sciences and School of Medical Sciences, The University of Sydney, Sydney, New South Wales 2006, Australia; Science and Technology Innovation Center, Shandong First Medical University & Shandong Academy of Medical Sciences, Taian 271000, China

**Keywords:** SARS-CoV-2, COVID-19, coronavirus, evolution, bats, phylogeny, spike protein, swine acute diarrhea syndrome, porcine epidemic diarrhea virus

## Abstract

Although a variety of SARS-CoV-2 related coronaviruses have been identified, the evolutionary origins of this virus remain elusive. We describe a meta-transcriptomic study of 411 samples collected from 23 bat species in a small (~1100 hectare) region in Yunnan province, China, from May 2019 to November 2020. We identified coronavirus contigs in 40 of 100 sequencing libraries, including seven representing SARS-CoV-2-like contigs. From these data we obtained 24 full-length coronavirus genomes, including four novel SARS-CoV-2 related and three SARS-CoV related genomes. Of these viruses, RpYN06 exhibited 94.5% sequence identity to SARS-CoV-2 across the whole genome and was the closest relative of SARS-CoV-2 in the ORF1ab, ORF7a, ORF8, N, and ORF10 genes. The other three SARS-CoV-2 related coronaviruses were nearly identical in sequence and clustered closely with a virus previously identified in pangolins from Guangxi, China, although with a genetically distinct spike gene sequence. We also identified 17 alphacoronavirus genomes, including those closely related to swine acute diarrhea syndrome virus and porcine epidemic diarrhea virus. Ecological modeling predicted the co-existence of up to 23 *Rhinolophus* bat species in Southeast Asia and southern China, with the largest contiguous hotspots extending from South Lao and Vietnam to southern China. Our study highlights both the remarkable diversity of bat viruses at the local scale and that relatives of SARS-CoV-2 and SARS-CoV circulate in wildlife species in a broad geographic region of Southeast Asia and southern China. These data will help guide surveillance efforts to determine the origins of SARS-CoV-2 and other pathogenic coronaviruses.

## Introduction

Most viral pathogens in humans have zoonotic origins, arising through occasional (e.g. coronavirus, Ebola virus) or frequent (e.g. avian influenza A virus) animal spillover infections. Bats (order Chiroptera) are the second most diverse mammalian order after Rodentia and currently comprise ~1420 species, accounting for some 22% of all named mammalian species (Letko et al., 2020). Bats are well known reservoir hosts for a variety of viruses that cause severe diseases in humans, and have been associated with the spillovers of Hendra virus, Marburg virus, Ebola virus and, most notably, coronaviruses. Aside from bats and humans, coronaviruses can infect a wide range of domestic and wild animals, including pigs, cattle, mice, cats, dogs, chickens, deer and hedgehogs (Chan et al., 2013; Su et al., 2016; Corman et al., 2018).

By 2019 there were six known human coronaviruses (HCoV): HCoV-229E, HCoV-OC43, severe acute respiratory syndrome coronavirus (SARS-CoV), HCoV-NL63, HCoV-HKU1, and Middle East respiratory coronavirus (MERS-CoV) (Su et al., 2016; Forni et al., 2017). HCoV-229E, HCoV-NL63, SARS-CoV and MERS-CoV were known to have zoonotic origins, with bats likely important reservoir hosts, although sometimes emergence in humans followed transmission through so-called “intermediate” hosts such as palm civets for SARS-CoV and dromedary camels for MERS-CoV (Corman et al., 2018; Ye et al., 2020). Similarly, it has been proposed that rodents may be the natural hosts of HCoV-OC43 and HCoV-HKU1, with cattle a possible intermediate host for HCoV-OC43 (Corman et al., 2018; Ye et al., 2020).

In early 2020, a novel coronavirus, SARS-CoV-2, was identified as the causative agent of a pneumonia outbreak in Wuhan, China, that eventually turned into a global pandemic (Zhu et al., 2020; Lu et al., 2020; Wu et al., 2020a). A combination of retrospective genome sequencing and ongoing sampling then identified a number of SARS-CoV-2 related coronaviruses in wildlife. These included: (i) the bat (*Rhinolophus affinis*) virus RaTG13 that is the closest relative of SARS-CoV-2 across the viral genome as a whole (Zhou et al., 2020b); (ii) the bat (*R. malayanus*) derived coronavirus RmYN02 that is the closest relative of SARS-CoV-2 in the long ORF1ab gene and which contains a similar nucleotide insertion at the S1/S2 cleavage site of the spike gene (Zhou H et al., 2020a); (iii) viruses from the Malayan pangolin (*Manis javanica*) that comprised two lineages reflecting their Chinese province of collection by local customs authorities (Guangdong and Guangxi), with the pangolins from Guangdong possessing identical amino acids at the six critical residues of the receptor binding domain (RBD) to human SARS-CoV-2 (Lam et al., 2020; Xiao et al., 2020); and (iv) a more distant SARS-CoV-2 related coronavirus from a bat (*R. cornutus*) sampled in Japan (Murakami et al., 2020). More recently, two novel betacoronaviruses (STT182 and STT200) were described in *R. shameli* bats sampled from Cambodia in 2010 that share 92.6% nucleotide identity with SARS-CoV-2 as well as five of the six critical RBD sites observed in SARS-CoV-2 (Hul et al., 2021). In addition, a novel bat (*R. acuminatus*) coronavirus isolated from Thailand (RacCS203) in June 2020 was recently identified and found to be closely related to RmYN02 (Wacharapluesadee et al., 2021). Collectively, these studies indicate that bats across a broad swathe of Asia harbor coronaviruses that are closely related to SARS-CoV-2 and that the phylogenetic and genomic diversity of these viruses has likely been underestimated. Herein, we report the discovery of additional bat coronaviruses from Yunnan province, China that reveal more of the diversity and complex evolutionary history of these viruses, including both cross-species transmission and genomic recombination.

## Results

### Identification of novel bat coronaviruses

From May 2019 to November 2020, a total of 283 fecal samples, 109 oral swabs and 19 urine samples were collected from bats in Yunnan province, China. The majority of samples were collected from horseshoe bats, comprising: *Rhinolophus malayanus* (n=88), *R. stheno* (n=36), *R. sinicus* (n=34), and *R. siamensis* (n=12), *R. pusillus* (n=2), other *Rhinolophus* sp. (n=11), and *Hipposideros larvatus* (n=59) (Figure 1A and 1B, Table S1). These samples were pooled into 100 libraries (numbered p1 to p100) according to the collection date and host species, with each library containing one to 11 samples. Meta-transcriptomic (i.e. total RNA) sequencing was performed and coronaviruses contigs were identified in 40 libraries (Table S2). Blastn searches of the *de novo* assemblies identified 26 long contigs (>23,000 nt in length) that mapped to coronavirus genomes present in 20 libraries, including nine sarbecoviruses (i.e. from the genus *Betacoronavirus)* and 17 alphacoronaviruses. The number of read-pairs mapping to these long contigs ranged from 3,433 to 21,498,614, with the average depth ranging from 35.86 to 215,065.00 (Table S3). It should be noted that pool p1 comprising 11 fecal samples from *R. malayanus* was the same pool previously used to identify RmYN01 and RmYN02 (Zhou et al., 2020a). The remaining 24 genomes were named in the same manner, in which the first two letters represent an abbreviation of the bat species, YN denotes Yunnan, and the final number is a serial number ranging from 03 to 26. In addition, several short contigs related to SARS-CoV-2 were identified in two other libraries - p7 and p11 (Figure S1, Table S2).

**Figure 1.**
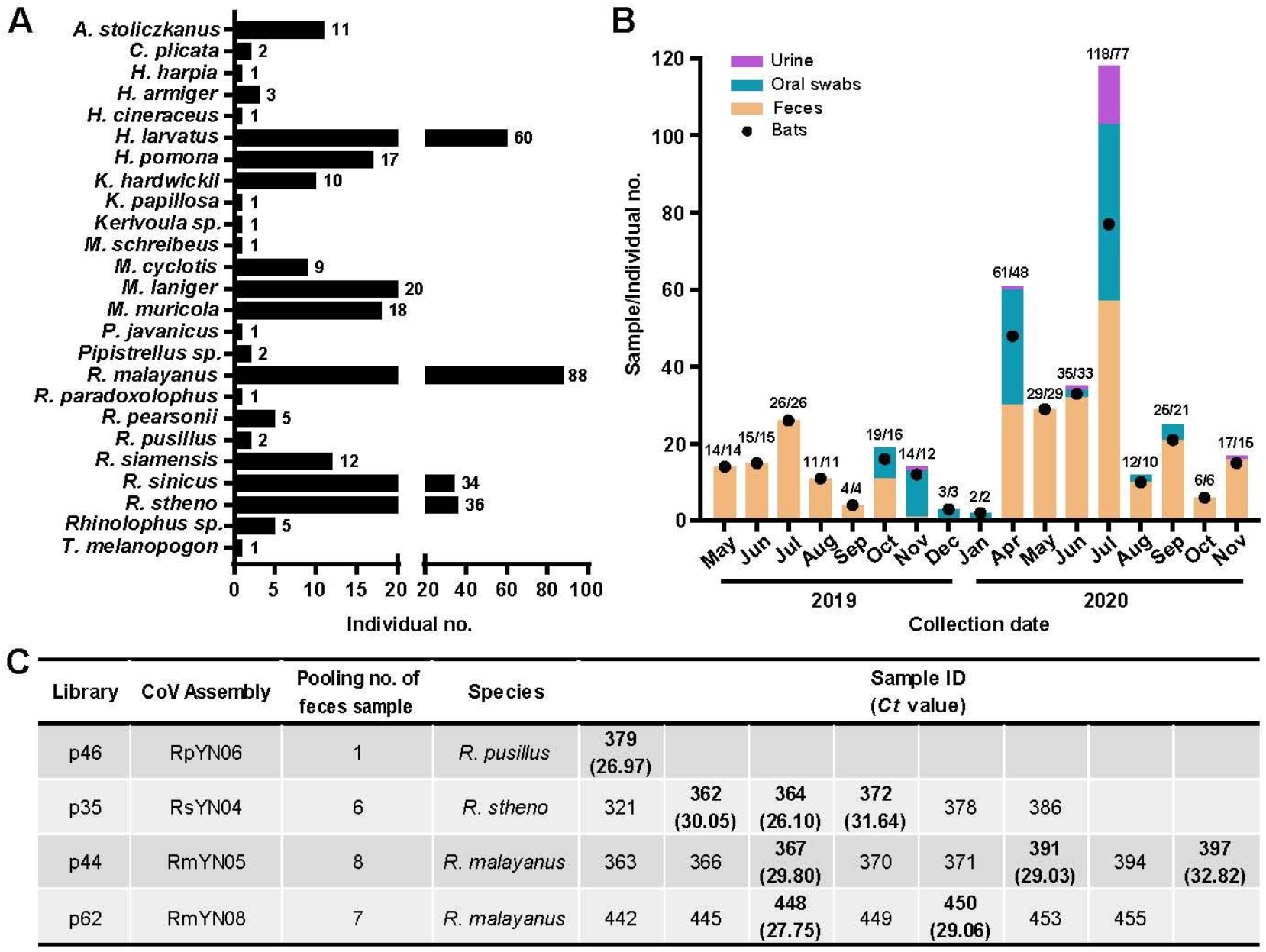
Sampling information and detection of SARS-CoV-2-like viruses in individual bat fecal samples. (A) Sample numbers of different bat species captured live in Yunnan province from May 2019 to November 2020. (B) Numbers of samples collected from different time points (orange column - feces; green - oral swab; light purple - urine). The numbers of individual bats are shown with black dots and relate to the y-axis. The associated numbers are in the form sample numbers/number of individual bats. (C) Identification of SARS-CoV-2-like virus positive samples using qPCR. Also see Tables S1 and S5.

Further Blastn analyses revealed that four of the seven novel sarbecoviruses identified here (RpYN06, RsYN04, RmYN05, and RmYN08) were related to SARS-CoV-2 with nucleotide identities ranging from 82.46% to 97.21%, while the remaining three (RsYN03, RmYN07, and RsYN09) were more closely related to SARS-CoV with nucleotide identities ranging from 91.60% to 93.28%. We next designed specific primers and a probe set of quantitative real-time PCR primers (qPCR) (Table S4) that targeted the conserved region of the1a gene region to detect the presence of the four SARS-CoV-2 related viruses in individual bats (i.e. prior to sample pooling; Figure 1C). Pool p46 only contained only a single positive fecal sample, no. 379, collected on May 25, 2020, and the virus was detected with a cycle threshold (*Ct*) value of 26.97 (Figure 1C). SARS-CoV-2 related virus was also detected in three (sample nos. 362, 364, and 372) of the six, three (sample nos. 367, 391, and 397) of the eight, and two (sample nos. 448 and 450) of the seven samples in pool nos. p35, 44 and 62, respectively, with *Ct* values ranging from 26.10 to 32.82 (Figure 1C). Among these, samples 362, 364, 372 and 367 were collected on May 25, 2020, 391 and 397 were collected on June 3, 2020, while both 448 and 450 were collected on July 16, 2020. The 5’ and 3’ termini and the spike gene sequences of the four coronaviruses related to SARS-CoV-2 were verified using individual samples 379, 364, 367 and 450 with 5’ and 3’ RACE (Table S4) and Sanger sequencing. Results from Sanger sequencing were consistent with those obtained from the meta-transcriptomic sequencing.

### Sequence identities between SARS-CoV-2 and related viruses

At the scale of the whole genome, RpYN06 exhibited 94.5% sequence identity to SARS-CoV-2, making it, after RaTG13 (96.0%), the second closest relative of SARS-CoV-2 documented to date (Figure 2). However, because of extensive recombination, patterns of sequence similarity vary markedly across the virus genome, and RmYN02 shared 97.18% sequence identity with SARS-CoV-2 in the 1ab open reading frame (ORF), compared to 97.19% for RpYN06. In addition to the ORF1ab, RpYN06 shared the highest nucleotide identities with SARS-CoV-2 in the RdRp (RNA-dependent RNA polymerase; 98.36%), ORF7a (96.72%), ORF8 (97.54%), N (97.70%), and ORF10 (100%) (Figure 2, Table S5 and S6). However, RpYN06 exhibited only 76.3% nucleotide identity to the SARS-CoV-2 spike gene and 60.9% in the receptor binding domain (RBD), thereby similar to RmYN02, ZC45, ZXC21 and the Thailand coronavirus strains (Figure 2). Excluding the spike gene, the sequence identities of RpYN06, RmYN02 and RaTG13 to SARS-CoV-2 were 97.17%, 96.41% and 96.49%, respectively.

**Figure 2.**
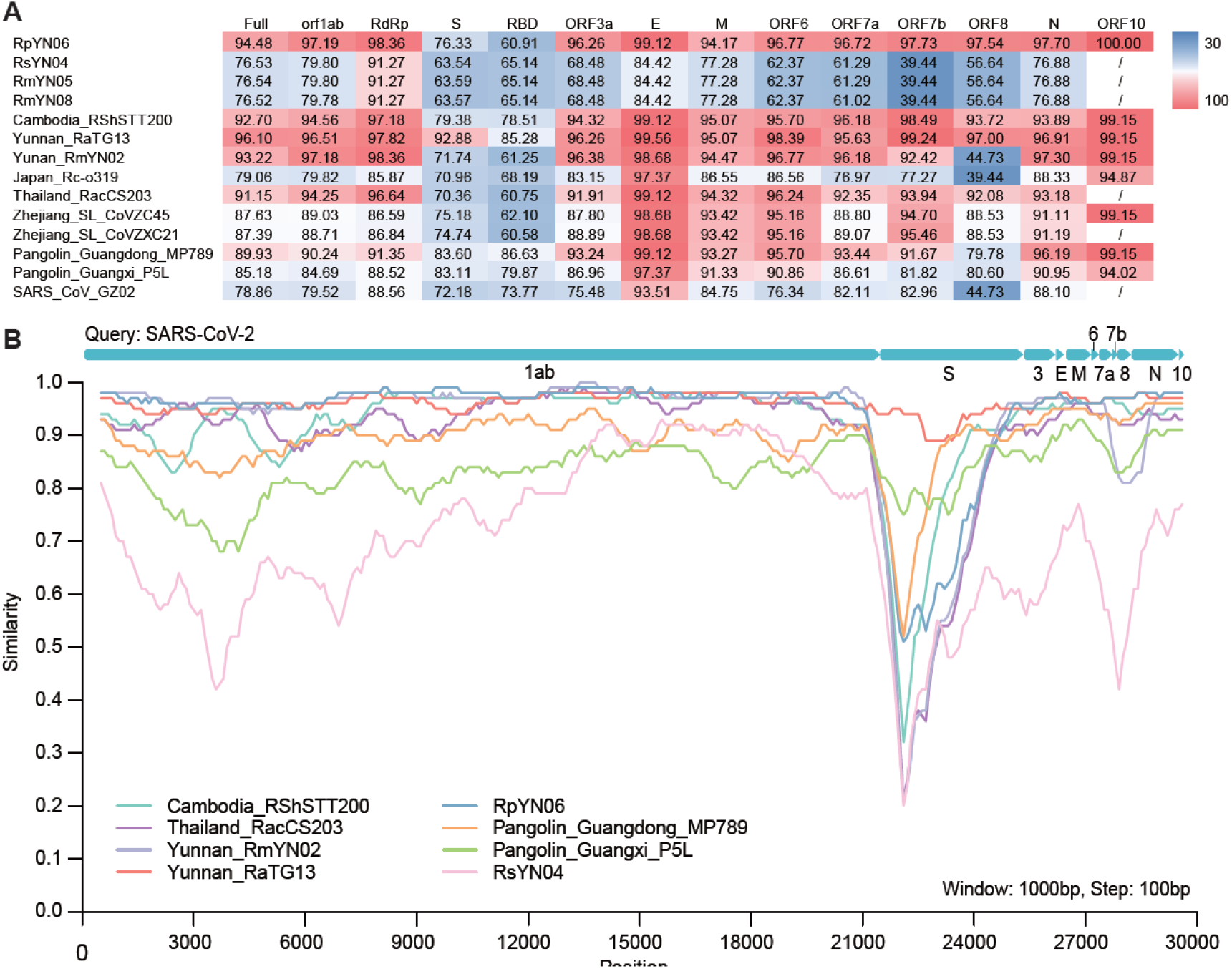
Sequence identities between SARS-CoV-2 and representative sarbecoviruses. (A) Pairwise sequence identities between SARS-CoV-2 (reference genome: NC_045512), and SARS-CoV-2 related coronaviruses. The degree of sequence similarity is highlighted by the shading, with cells shaded red denoting the highest identities. (B) Whole genome sequence similarity plot of nine SARS-CoV-2 related coronaviruses using the SARS-CoV-2 as a query. The analysis was performed using Simplot, with a window size of 1000bp and a step size of 100bp. Also see Tables S2 and S6.

In contrast, RsYN04, RmYN05 and RmYN08 exhibited >99.96% nucleotide identities to each other at the scale of the whole genome. Such strong similarity is indicative of viruses from the same species, even though they were sequenced on different lanes and the samples were collected from different bat species at different time points. In addition, they shared low nucleotide identities with SARS-CoV-2 across the whole genome (76.5%), particularly in the spike gene, ORF3a, ORF6, ORF7a, ORF7b and ORF8 with nucleotide identities <70% (Figure 2). Interestingly, when using RsYN04 as the query sequence, the closest hit in the Blastn search was the pangolin derived coronavirus MP789 (MT121216.1) with 82.9% nucleotide identity.

### Evolutionary history of sarbecoviruses

Phylogenetic analysis of full-length genome sequences of representative sarbecoviruses revealed that SARS-CoV-2 was most closely related to RaTG13, while RmYN02 and the Thailand strains formed a slightly more divergent clade. Notably, RpYN06 was placed at the basal position of the clade containing SARS-CoV-2 and its closest relatives from bats and pangolins (Figure 3A, Table S6). In contrast, RsYN04, RmYN05 and RmYN08 grouped together and clustered with the pangolin derived viruses from Guangxi, although being separated from them by a relatively long branch. Finally, three SARS-CoV related coronaviruses (RsYN03, RmYN07, and RsYN09) fell within the SARS-CoV lineage, grouping with other bat viruses previously sampled in Yunnan (Figure 3A).

**Figure 3.**
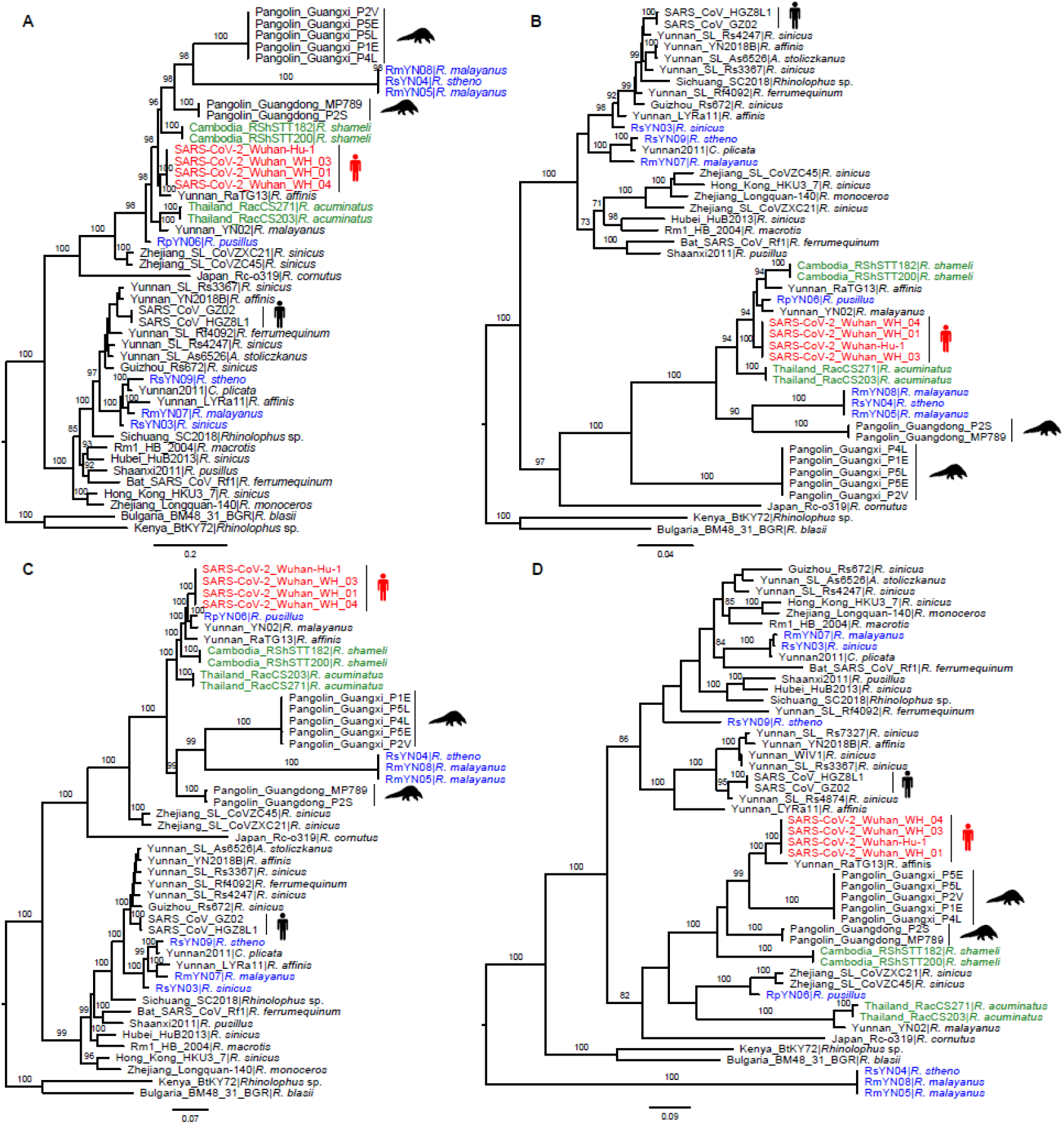
Phylogenetic analysis of SARS-CoV-2 and representative sarbecoviruses. Nucleotide sequence phylogenetic trees of (A) the full-length virus genome, (B) the RdRp gene, (C) the ORF1ab, and (D) the spike gene. The phylogenetic trees in panels A-C were rooted using the bat viruses Kenya_BtKY72 (KY352407) and Bulgaria_BM48_31_BGR (GU190215) as outgroups, whereas the tree in panel D was midpoint rooted for clarity only. Phylogenetic analysis was performed using RAxML (Stamatakis 2014) with 1000 bootstrap replicates, employing the GTR nucleotide substitution model. Branch lengths are scaled according to the number of nucleotide substitutions per site. Viruses are color-coded as follows: red - SARS-CoV-2; blue - new genomes generated in this study; green - recently published sequences from Thailand and Cambodia. Also see Table S6.

A different topological pattern was observed in the phylogeny of the RdRp (Figure 3B). In particular, RpYN06 now grouped with RmYN02 (although with weak bootstrap support), with together formed a clade with RaTG13, the two Cambodian strains, and SARS-CoV-2 (Figure 3B). The two bat derived strains from Thailand formed a separate lineage. Perhaps more striking was that RsYN04, RmYN05 and RmYN08 now grouped with the Guangdong pangolin viruses (rather than those from Guangxi; Figure 3B). A different pattern again was observed in the phylogeny of the entire ORF1ab (Figure 3C). RpYN06 and RmYN02 now formed a clade and that was the direct sister-group to SARS-CoV-2, with RaTG13 a little more divergent (Figure 3C). In addition, RsYN04, RmYN05 and RmYN08 now clustered with the pangolin derived strains from Guangxi (Figure 3C), consistent with the complete genome phylogeny.

In the spike gene phylogeny, SARS-CoV-2 and RaTG13 still grouped together, with both pangolin lineages falling as sister groups (Figure 3D). The two Cambodian bat viruses formed a separate and more divergent lineage. Strikingly, RpYN06 exhibited marked phylogenetic movement, this time clustering with two previously described bat viruses from Zhejiang province - ZC45 and ZXC21 - whereas the Thailand bat virus clustered closely with RmYN02 (Figure 3D). In addition, RsYN321B, RmYN363B, and RmYN442B did not fall within the SARS-CoV and SARS-CoV-2 clades, but instead formed a separate and far more divergent lineage (Figure 3D). Finally, in the phylogeny of the RBD region, SARS-CoV-2 clustered with the pangolin viruses from Guangdong with the two Cambodian bat viruses the next most closely related viruses (Figure S2). RpYN06 fell within a lineage comprising several bat derived betacoronaviruses, including ZC45, ZXC21, RsYN09, RsYN03, and RmYN07. As expected given the complete S gene tree, bat viruses RsYN04, RmYN05 and RmYN08 grouped together and formed a lineage, characterized by a long branch (Figure S2).

### Molecular characterizations of the spike protein of the novel bat sarbecoviruses

At the six amino acid positions deemed critical for binding to the human angiotensin-converting enzyme 2 (hACE2) receptor, SARS-CoV-2 and the three bat derived viruses identified here (RsYN04, RmYN05 and RmYN08) shared L455 and Y505. In contrast, despite being a closer overall relative, RpYN06 only possessed one identical amino acid with SARS-CoV-2 - Y505 (Figure 4A). In the S1/S2 cleavage site of the spike gene, none of the four SARS-CoV-2 related viruses reported here possessed a similar insertion/deletion (indel) pattern as SARS-CoV-2 (Garry et al., 2021) (Figure 4A). Interestingly, however, the recently sampled bat virus from Thailand possessed a PVA three amino acid insertion at this site, similar to the PAA insertion found in RmYN02. In addition, two indel events have been identified in the RBD of many bat associated coronaviruses (Holmes et al., 2021), including RpYN06 that was characterized by indel patterns identical to those of ZC45 and ZXC21 (Figure 4A). There were no indel events in SARS-CoV-2 and the pangolin derived coronaviruses in RBD, and RsYN04, RmYN05 and RmYN08 possessed one unique indel event different from other sarbecoviruses (Figure 4A). In addition, and similar to other bat derived coronaviruses, the four novel SARS-CoV-2 related viruses possessed several indel events in the N-terminal domain, while RsYN04, RmYN05 and RmYN08 again possessed a unique indel pattern (Figure S3). Notably, RpYN06, ZC45, ZXC21 and the Guangdong pangolin virus shared the same indel pattern, with RpYN06 exhibiting high amino acid identity to these viruses in the N-terminal domain (amino acid identities ranging from 85.3% to 99.0%; Figure S4).

**Figure 4.**
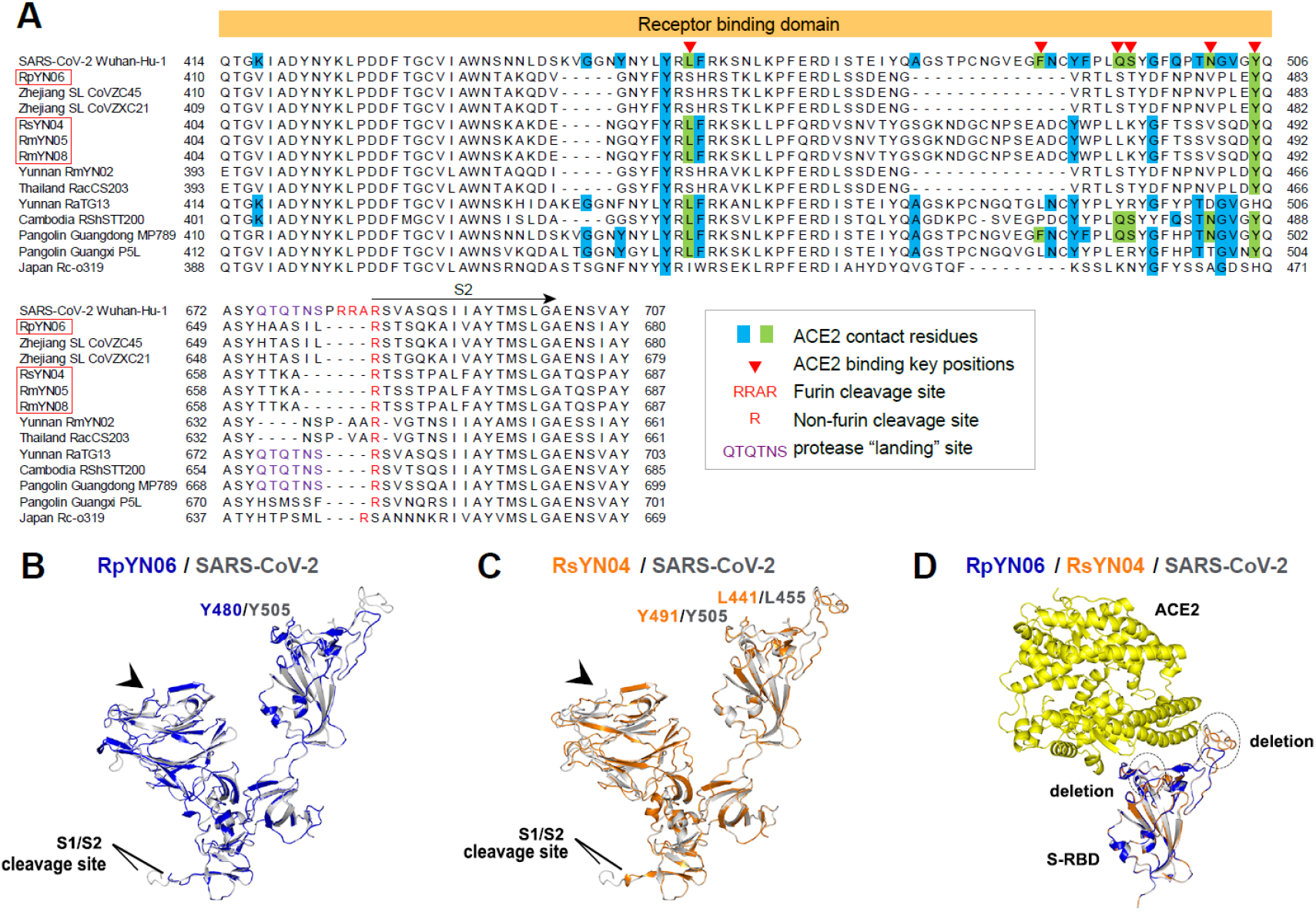
Molecular characterizations of the RBD and homology modeling of the S1 subunit of the novel sarbecoviruses. (A) Sequence alignment of the RBD region of SARS-CoV-2 and representative betacoronavirus genomes (annotation following Holmes et al., 2021). (B-C) Homology modeling and structural comparison of the S1 subunit between (B) RpYN06 and SARS-CoV-2, and (C) RsYN04 and SARS-CoV-2. (D) Structural similarity between the RpYN06:hACE2, RsYN04:hACE2 and SARS-CoV-2-RBD:hACE2 complexes. The three-dimensional structures of the S1 from RpYN06, RsYN04 and SARS-CoV-2 were modeled using the Swiss-Model program (Waterhouse et al., 2018) employing PDB: 7A94.1 as the template. The S1 domains of RpYN06, RsYN04 and SARS-CoV-2 are colored blue, orange and gray, respectively. The hACE2 are colored yellow. The deletions in RpYN06 and/or RsYN04 are highlighted. The NTD (black arrow heads) is marked. Also see Figure S3.

We predicted and compared the three-dimensional structures of RpYN06, RsYN04 and SARS-CoV-2 using homology modeling (Figures 4B–4D). In a similar manner to RmYN02 (Zhou et al., 2020a), the RBD of RpYN06 had two shorter loops than those observed in SARS-CoV-2, while RsYN04 only had one shorter loop (Figure 4D). In addition, near the S1/S2 cleavage sites, the conformational loop of RpYN06 and RsYN04 were different from those of SARS-CoV-2 (Figures 4B–4C). Notably, RsYN04 exhibited greater amino acid identity (71.28%) and shared more structural similarity with the SARS-CoV-2 RBD than RpYN06 (63.08%). Importantly, the conformational variations caused by these amino acid substitutions and deletions were speculated to interfere with the binding of RpYN06 and RsYN04 RBD to hACE2 (Figure 4D). However, RsYN04 exhibited lower structural similarity with SARS-CoV-2 in the N-terminal domain (NTD) (Figure 4C, black arrowheads, 39.19% amino acid identity) than RpYN06 (65.87% amino acid identity).

### Phylogenetic analysis of the novel bat alphacoronaviruses

As well as betacoronaviruses, we identified 17 novel bat alphacoronaviruses. Phylogenetic analyses of the full-length genomes (Figure 5A), the RdRp genes (Figure 5B), and ORF1ab (Figure S5) of these 17 alphacoronaviruses and representative background viruses were consistent, with all trees revealing that the viruses newly identified here fell within four established subgenera: *Decacovirus* (n=12), *Pedacovirus* (n=1), *Myotacovirus* (n=1), and *Rhinacovirus* (n=2) (Figure 5). Of particular note were MlYN15 and RsYN25 isolated from *Myotis laniger* and *R. stheno* bats that were closely related to swine acute diarrhea syndrome coronavirus (SADS-CoV) (Figure 5) (Zhou et al., 2018) sharing nucleotide identities 87.55% - 87.61%. In addition, HlYN18, isolated from a *Hipposideros larvatus* bat, fell within the subgenus *Pedacovirus*, and was close to the porcine epidemic diarrhea virus (PEDV) lineage (Figure 5). Notably, the virus CpYN11 (isolated from *Chaerephon plicatus)* clustered with WA3607 (GenBank accession no. MK472070; isolated from a bat from Australia), which together might represent an unclassified subgenus (Figure 5). Finally, RsYN14, RmYN17, McYN19, and RmYN24, although isolated from different bat species and sequenced on different lanes, they were almost identical (with nucleotide identity >99.98% to each other) and might represent a novel species of subgenus *Decacovirus*.

**Figure 5.**
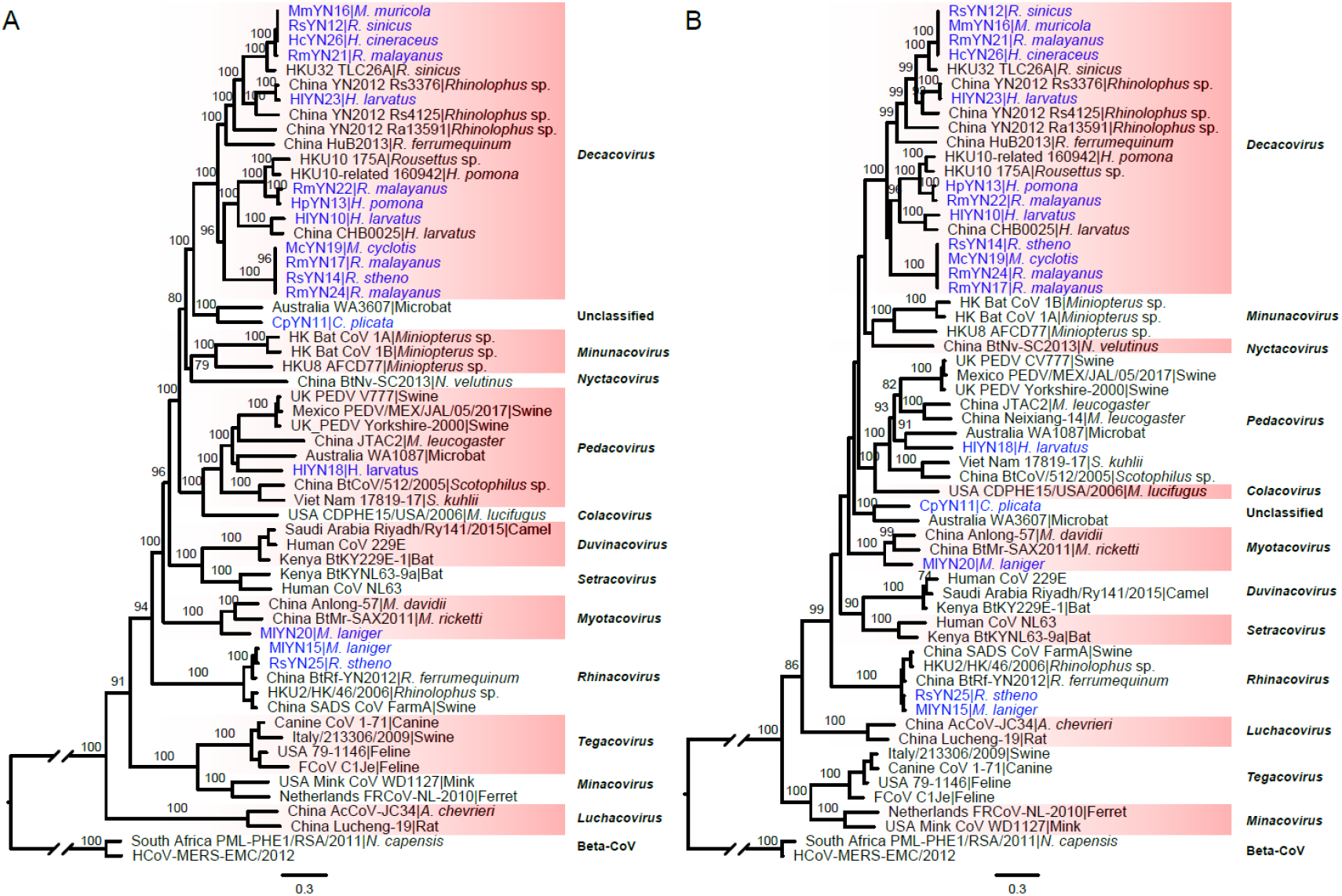
Phylogenetic analysis of 17 novel alphacoronaviruses and representative viruses from different subgenera. Phylogenetic trees of (A) the full-length virus genome and (B) the RdRp gene of alphacoronaviruses. Phylogenetic analysis was performed using RAxML(Stamatakis 2014) with 1000 bootstrap replicates, employing the GTR nucleotide substitution model. The two trees were rooted using two betacoronaviruses as outgroups - South_Africa_PML-PHE1/RSA/2011 (KC869678.4) and HCoV-MERS-EMC (NC_019843). Branch lengths are scaled according to the number of substitutions per site. Also see Figures S4 and S8.

Although the phylogenetic trees of the spike gene (Figure S6A) and protein sequences (Figure S6B) were topologically similar to those of the full-length genome, RdRp and ORF1ab, a number of notable differences were apparent indicative of past recombination events. First, CpYN11 clustered with HKU8 rather than WA3607 in the spike gene tree where they formed a separate lineage. Second, the topology of the subgenus *Decacovirus* in the spike gene tree was different to those observed in other gene regions. Finally, the two viruses belonging to the subgenus *Tegacovirus* were placed into the subgenera *Pedacovirus* (GenBank accession no. NC_028806) and a separate lineage (GenBank accession no. DQ848678), respectively.

### Ecological modeling of the distribution of *Rhinolophus* species in Asia

To better understand the ecology of bat coronaviruses, we modeled the distribution of 49 *Rhinolophus* species in Asia using the collated distribution data and several ecological measures (Figures 6 and S7). The models performed well with a mean Area Under Curve (AUC) of 0.96 for training and 0.92 for testing, and all training AUCs were above 0.88. Continentality (reflecting the difference between continental and marine climates) was, on average, the most important factor, contributing an average of 14.91% (based on permutation importance), followed by temperature seasonality at 11.7% average contribution, mean diurnal temperature range at 5.69%, and annual potential evapotranspiration at 5.38%. Three additional ecological factors also contributed over 5% on average: minimum precipitation at 5.25%, potential evapotranspiration seasonality at 5.17% and Emberger’s pluviothermic quotient (a measure of climate type) at 5%. The next most important factor was the distance to bedrock (an indicator of potential caves and rock outcrops) at 4.46%. Thus, local climate, especially factors that influence diet availability across the year, is seemingly key to determining bat species distributions across the region.

**Figure 6.**
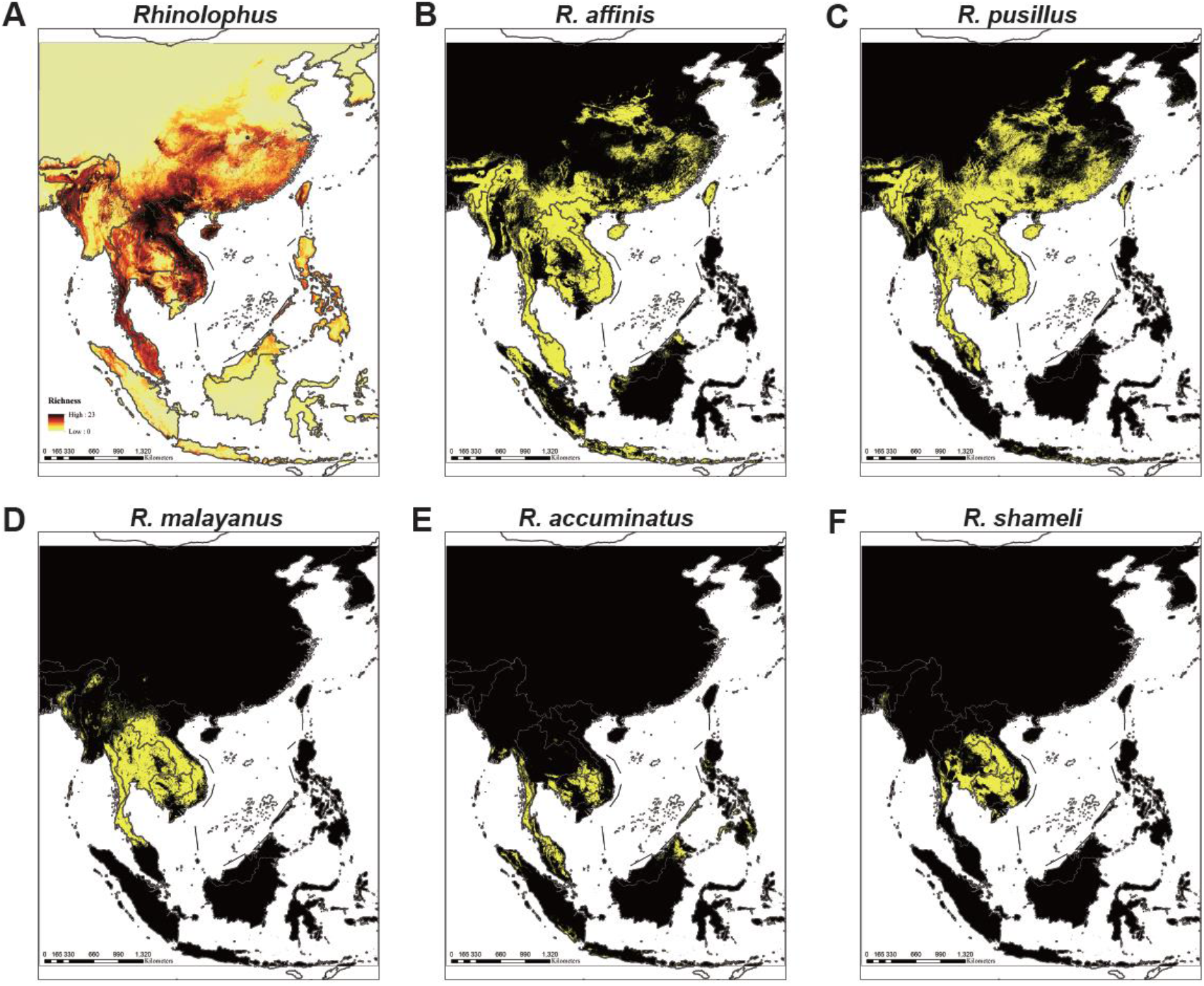
Ecological modeling the geographical distribution of 49 Rhinolophid bat species. (A) Models of 49 *Rhinolophus* bat species that predict diversity in five regions covering mainland Southeast Asia, Philippines, Java-Sumatra, Borneo and Sulawesi-Moluccas. The map color represents species richness, with up to 23 species projected to co-exist. (B-F) Location distribution of (B) the RaTG13 host species *R. affinis*, (C) the RpYN06 host species *R. pusillus*, (D) the RmYN02 host species *R. malayanus*, (E) the RacCS203 host species *R. accuminatus*, and (F) the STT182 and STT200 host species *R. shameli*. The yellow region represents the predicted range of each species. Also see Figure S7.

Although we could not accurately model diversity for Indonesia because of limited recently available data and likely high endemism, mainland Southeast Asia was well mapped (Figures 6 and S7). Most of mainland Southeast Asia’s remaining tropical forests showed a high diversity of Rhinolophid bats, with a maximum of 23 species estimated to exist concurrently (Figure 6A). Rhinolophid hotspots occurred in forests throughout much of mainland Southeast Asia, with the largest contiguous hotspots extending from South Lao and Vietnam to Southern China (Figure 6A). Hotspots were also identified in the Hengduan mountains, and some parts of northern Myanmar and Nagaland in India (Figure 6A).

Interestingly, *R. affinis* (Figure 6B) and *R. pusillus* (Figure 6C) were widely distributed in Southeast Asia and southern China, and most bat species shared hotspots in Cambodia and peninsula Thailand. Several Rhinolophid species extended their ranges northwards into southern China reflecting the presence of forest (*R. affinis* and *R. pusillus*), whereas the geographic range of *R. malayanus* only just reached southern China (Figures 6D–6F). Ecological drivers for these species unsurprisingly showed some differences. Specifically, *R. affinis* was also influenced by temperature seasonality (16.59%), followed by Emberger’s pluviothermic quotient and mean diurnal range (8.79, and 8.7%), while *R. malayanus* (a smaller species) was mainly influenced by annual potential evapotranspiration mean (33.79%) and seasonality (14.57%). *R. pusillus* was influenced by temperature seasonality (12.44%) and continentality (9%), and *R. shameli* was largely influenced by annual potential evapotranspiration seasonality (34.81%) followed by annual evapotranspiration (9.79%). Overall, these factors control the range limits, and food availability for these bat species.

It should be noted that the ecological modeling identified several other Rhinolophid species with wide geographic distributions: *R. huanensis, R. lepidus, R. luctus, R. macrotis, R. marshalli, R. microglobosus, R. pearsoni, R. rouxii, R. stheno, R. thomasi*, and *R. yunnanensis* (Figure S7). Notably, *R. stheno* was found to host both SARS-CoV-2 and SARS-CoV-like coronaviruses in the present study.

## Discussion

To reveal more of the diversity, ecology and evolution of bat viruses, we collected bat samples in Yunnan province, China during 2019-2020. Overall, 40 of the 100 sequencing libraries contained coronaviruses, including seven libraries with contigs that could be mapped to SARS-CoV-2. In particular, we assembled 24 novel coronavirus genomes from different bat species, including four SARS-CoV-2 like coronaviruses. Further PCR based tests revealed that these four viruses tested positive in nine individual samples collected in Yunnan province between May and July 2020. Together with the SARS-CoV-2 related virus collected from Thailand in June 2020 (Wacharapluesadee et al., 2021), these results clearly demonstrate that SARS-CoV-2 related viruses continue to circulate in bat populations, and in some regions the prevalence of SARS-CoV-2 related coronaviruses might be relatively high.

Of particular note was that one of the novel bat coronavirus identified here - RpYN06 - exhibited 94.5% sequence identity to SARS-CoV-2 across the genome as a whole and in some individual gene regions (ORF1ab, ORF7a, ORF8, N, and ORF10) was the closest relative of SARS-CoV-2 identified to date. However, the low sequence identity in the spike gene, itself clearly the product of a past recombination event, made it a second closest relative of SARS-CoV-2, next to RaTG13, at the genomic scale. Hence, aside from the spike gene, RpYN06 possessed a genomic backbone that is arguably the closest to SARS-CoV-2 identified to date.

Although several SARS-CoV-2-like viruses have been identified from different wildlife species that display high sequence similarity to SARS-CoV-2 in some genomic regions, none are highly similar (e.g. >95%) to SARS-CoV-2 in the spike gene in terms of both the overall sequence identity and the amino acid residues at critical receptor binding sites (Zhou et al., 2020b; Lam et al., 2020; Xiao et al., 2020; Zhou et al., 2020a; Murakami et al., 2020; Hul et al., 2021; Wacharapluesadee et al., 2021). Indeed, the spike protein sequences of three of the novel coronaviruses described here (RsYN04, RmYN05, RmYN08) formed an independent lineage separated from known sarbecoviruses by a relatively long branch. In this context it is interesting that the recently identified bat coronavirus from Thailand carried by a three-amino acid-insertion (PVA) at the S1/S2 cleavage site (Wacharapluesadee et al., 2021). Although this motif is different to that seen in SARS-CoV-2 (PRRA) and RmYN02 (PAA), this once again reveals the frequent occurrence of indel events in the spike proteins of naturally sampled betacoronaviruses (Garry et al., 2021; Holmes et al., 2021). Collectively, these results highlight the extremely high, and likely underestimated, genetic diversity of the sarbecovirus spike proteins, and which likely reflects their adaptive flexibility.

Previous studies have revealed frequent host switching of coronaviruses among bats (Latinne et al., 2020). Indeed, we identified nearly 100% identical coronaviruses from multiple different bat species both in *Alphacoronavirus* and *Betacoronavirus*, indicative of the frequent cross-species virus transmission that drives virus evolution. This in part likely reflects their roosting behavior and propensity to share the same or close habitats. However, while facilitating host jumping, that individual bat species can harbor multiple viruses increases the difficulty in resolving the origins of SARS-CoV-2 and other pathogenic coronaviruses. Of particular note was that the three of the newly identified SARS-CoV-2 like coronaviruses grouped together with the pangolin derived coronaviruses from Guangxi in the whole genome phylogeny. Although the associated branch lengths are relatively long such that other hosts may be involved, and there are some topological differences between gene trees, this is suggestive of virus transmission between pangolins and bats. Recently, a new SARS-CoV-2 related coronavirus was identified from a pangolin from Yunnan (GISAID ID EPI_ISL_610156). Whether pangolin derived coronaviruses have formed a separate lineage clearly warrants further investigation.

Rhinolophid bats are important hosts for coronaviruses (Fan et al., 2019; Latinne et al., 2020). Our ecological modeling revealed high richness of Rhinolophids across much of Southeast Asia and southern China, with up to 23 species projected to co-exist from the 49 species included in analysis. The largest expanses of high bat diversity habitat stretch from South Vietnam into southern China (Hughes et al., 2012; Allen et al., 2017). Indeed, it is striking that all the bat viruses described here, as well as RmYN01 and RmYN02 described previously (Zhou et al., 2020a), were identified in a small area (~1100 hectare) in Yunnan province. This highlights the remarkable phylogenetic and genomic diversity of bat coronaviruses in a tiny geographic area and to which humans may be routinely exposed. Importantly, in addition to Rhinolophids, this broad geographic region in Asia is rich in many other bat families (Anthony et al., 2017) and other wildlife species (Olival et al., 2017) that have been shown to be susceptible to SARS-CoV-2 *in vitro* (Conceição et al., 2020; Wu et al., 2020; Sang et al., 2020; Yan et al., 2021). It is therefore essential that further surveillance efforts should cover a broader range of wild animals in this region to help track ongoing spillovers of SARS-CoV-2, SARS-CoV and other pathogenic viruses from animals to humans.

## Acknowledgments

This work was supported by the Academic Promotion Programme of Shandong First Medical University (2019QL006), the Key research and development project of Shandong province (2020SFXGFY01 and 2020SFXGFY08), the National Science and Technology Major Project (2020YFC0840800 and 2018ZX10101004-002), the National Major Project for Control and Prevention of Infectious Disease in China (2017ZX10104001-006), the Strategic Priority Research Programme of the Chinese Academy of Sciences (XDB29010102 and XDA20050202), the Chinese National Natural Science Foundation (32041010 and U1602265), and the High-End Foreign Experts Program of Yunnan Province (Y9YN021B01). W.S. was supported by the Taishan Scholars Programme of Shandong Province (ts201511056). Y.B. is supported by the NSFC Outstanding Young Scholars (31822055) and Youth Innovation Promotion Association of CAS (2017122). E.C.H. is supported by an ARC Australian Laureate Fellowship (FL170100022). We thank all the scientists who kindly shared their genomic sequences of the coronaviruses used in this study.

## Author Contributions

W.S., E.C.H. and A.C.H. designed and supervised research. X.C., Y.C. and A.C.H. collected the samples. H.Z., Y.B., M.C. and Y.Z. processed the samples. H.Z. performed the 5’ and 3’ RACE, Sanger sequencing and molecular detection. J.J., J.L. and T.H. performed the genome assembly and annotation. J.J., H.Z. and J.L. performed the genome analysis and interpretation. J.L. and H.S. performed the homology modelling. A.C.H. performed the ecological modeling. X.C. and Y.B. assisted in data interpretation and edited the paper. W.S., E.C.H. and A.C.H. wrote the paper.

## Declaration of Competing Interests

The authors declare no competing interests.

## STAR Methods

### RESOURCE AVAILABILITY

#### Lead Contact

Further information and requests for resources and reagents should be directed to and will be fulfilled by the Lead Contact, Weifeng Shi.

#### Materials Availability

This study did not generate new unique reagents.

### EXPERIMENTAL MODEL AND SUBJECT DETAILS

A total of 23 different bat species were tested in this study (Table S1). Samples were collected between May 2019 and November 2020 from Mengla County, Yunnan Province in southern China (101.271563 E, 21.918897 N; 101.220091 E, 21.593202 N and 101.297471 E, 21.920934 N). The Xishuangbanna Tropical Botanical Garden has an ethics committee that provided permission for trapping and bat surveys within this study.

### METHOD DETAILS

#### Sample collection

A total of 411 samples from 342 bats were collected from the Xishuangbanna Tropical Botanical Garden and its adjacent areas, Mengla County, Yunnan Province in southern China between May 2019 and November 2020. Bats were trapped using harp traps and a variety of samples were collected from each individual bat including feces (n=283), oral swab (n=109) and urine (n=19). Fecal and swab samples were collected and stored in RNAlater (Invitrogen), and urine samples were directly collected in the RNase-free tubes. These bats were primarily identified according to morphological criteria and found to belong to 23 different species, with the majority representing horseshoe bats (n=183) containing *Rhinolophus malayanus, R. stheno, R. sinicus, R. siamensis, R. pusillus* and other *R*. genus bats, as well as *Hipposideros larvatus* (n=59) (Table S1). All bats were sampled alive and subsequently released. All samples were transported on ice and then kept at −80°C until use.

#### Next generation sequencing

All bat samples were merged into 100 pools to generate sequencing libraries, based on the sample types, bat species and collection date. Of these bat libraries, 18 libraries have been described previously (Zhou et al., 2020a), including the library from which the viruses RmYN01 and RmYN02 were identified. These 18 libraries were combined with 82 additional libraries newly obtained here. Total RNA from samples was extracted using RNAprep pure Cell/Bacteria Kit (TianGen) and aliquots of the RNA solutions were then pooled in equal volume. Libraries were constructed using the NEBNext Ultra Directional RNA Library Prep Kit for Illumina (NEB). Ribosomal (r) RNA of fecal, oral swab and urine was removed using the TransNGS rRNA Depletion (Bacteria) Kit (TransGen) and rRNA of tissues was removed using TransNGS rRNA Depletion (Human/Mouse/Rat) Kit (TransGen), respectively. Paired-end (150 bp) sequencing of each RNA library was performed on the NovaSeq 6000 platform (Illumina) with the S4 Reagent Kit, and performed by the Novogene Bioinformatics Technology (Beijing, China).

#### Genome assembly and annotation

Clean reads from the next generation sequencing were classified with Kraken (v2.0.9) based on all microbial sequences from the NCBI nucleotide database. Paired-end reads classified as from coronaviruses were extracted from the Kraken output. To further verify the existence of coronaviruses, reads classified as coronaviruses were assembled with MEGAHIT (v1.2.9). The contigs from MEGAHIT were searched by BLASTn based on the NCBI nt database. Sequencing libraries with contigs identified as representing coronavirus were *de novo* assembled with coronaSPAdes (v3.15.0). The near complete genomes of coronavirus were then identified from the results of coronaSPAdes by BLASTn searching.

The newly assembled coronavirus genomes were validated by read mapping using Bowtie2 (v2.4.1). The coverage and depth of coronavirus genomes were calculated with SAMtools (v1.10) based on SAM files from Bowtie2. To further improve the quality of the genome annotations, SAM files of the reads mapping to SARS-CoV-2 were checked manually with Geneious (v2021.0.1), extending the ends as much as possible. The open reading frames (ORFs) of the verified genome sequences were annotated using Geneious (v2021.0.1) and then checked with closed references from NCBI. The taxonomy of these newly assembled coronavirus genome were determined by online BLAST (https://blast.ncbi.nlm.nih.gov/Blast.cgi).

Coronavirus contigs produced by MEGAHIT (v1.2.9) were analyzed to evaluate the existence of coronavirus sequences in each library. To mitigate the possibility of false positives due to index hopping, coronavirus contigs from different libraries within the same chip and same lane were compared, and if a shorter contig shared >99% nucleotide sequence identity with a longer contig from another library, the shorter one was removed.

#### Sanger sequencing

The assembled genome sequences of the beta-CoVs identified here were further confirmed by quantitative real-time PCR (qPCR), PCR amplification and Sanger sequencing. A TaqMan-based qPCR was first performed to test the feces of pools p19, p35, p44, p46, p52 and p62, as these contained beta-CoVs according to the metagenomic analysis. cDNA synthesis was performed using the ReverTra Ace qPCR RT Kit (TOYOBO). The qPCR reaction was undertaken using a set of probe and primer pairs (Table S4) in the *Pro Taq* HS Premix Probe qPCR Kit (AG) with a LightCycler 96 Real-Time PCR System (Roche).

##### Rapid amplification of cDNA ends (RACE)

The sequences of the 5’ and 3’ termini were obtained by RACE using the SMARTer RACE 5’/3’ Kit and 3’-Full RACE Core Set (Takara), according to the manufacturer’s instructions with some minor modifications. Two sets of gene-specific primers (GSPs) and nested-GSPs (NGSPs) for 5’ and one set for 3’ RACE PCR amplification were designed based on the assembled genome sequences of six beta-CoVs (Table S3). The first round of amplification was performed by touchdown PCR, while the second round comprised regular PCR. The PCR amplicons of ~1000 bp fragments of the two regions were obtained separately and sequenced with the amplified primer or gel purified followed by ligation with the pMD18-T Simple Vector (TaKaRa) and transformation into competent *Escherichia coli* DH5α (Takara). Insertion products were sequenced with M13 forward and reverse primers.

##### Amplification of beta-CoVs S gene and the host COI gene

Based on the spike gene and the adjacent sequences of RsYN04, RmYN05, RmYN08 and RpYN06, 9 primer pairs were designed for Sanger sequencing (Table S4). The cDNAs reverse transcribed above were used as templates. The thermal cycling parameters of PCR amplification were as follows: 5 mins at 95°C; followed by 30 s at 95°C, 30 s at 50°C (an exception of 55°C for primers 379SF5/379SR5), 1 min at 72°C for 30 cycles; and 10 min at 72°C. A second round PCR was then performed under the same conditions with the corresponding PCR products used as templates. Further confirmation of host species was based on analysis of the cytochrome b (*cytb*) gene obtained from the assembled contigs. We also amplified and sequenced the fragment of cytochrome c oxidase subunit I (*COI*) gene using primers VF1/VR1 (Ivanova et al., 2007). Briefly, the following touchdown PCR conditions were used: 30 s at 95°C, 30 s at 52°C to 45°C, 45 s at 72°C for 14 cycles; and followed by 30 s at 95°C, 30 s at 45°C, 45 s at 72°C for 30 cycles.

#### Bioinformatics analyses

##### Phylogenetic analysis

Multiple sequence alignment of the alphacoronavirus and betacoronavirus nucleotide sequences was performed using MAFFT (v7.450). Phylogenetic analysis of the complete genome and major genes were performed using the maximum likelihood (ML) method available in RAxML (v8.2.11) with 1000 bootstrap replicates, employing the GTR nucleotide substitution model and a gamma distribution of rate variation among sites. The resulting phylogenetic trees were visualized using Figtree (v1.4.4).

##### Sequence identity and recombination analysis

Pairwise sequence identity of the complete viral genome and genes between SARS-CoV-2 and representative sarbecoviruses was calculated using Geneious (v2021.0.1). A whole genome sequence similarity plot was performed using Simplot (v3.5.1), with a window size of 1000bp and a step size of 100bp.

##### Site and structural analysis of the spike gene

The three-dimensional structures of the S1 protein from RpYN06, RsYN04 and SARS-CoV-2 were modeled using the Swiss-Model program (Waterhouse et al., 2018) employing PDB: 7A94.1 as the template. Molecular images were generated with an open-source program - PyMOL. Multiple sequence alignment of spike gene amino acid sequences was performed with Clustal Omega (v1.2.2).

#### Ecological modeling

Data was collated using a combination of that from Hughes 2019, from various online repositories (Table S7), and additional GBIF data collated between 2017 and 2021. Further data was downloaded for Indonesia since 1990, even though wide-scale deforestation means that most species are to still likely to occupy small parts of their range. This provided sufficient data to model 49 Rhinolophid species based on 8418 occurrence points (once any duplicate points of species recorded repeatedly at the same location had been removed), with almost all records collected since 1998. Variables were selected to provide a good simulation of the environmental conditions that may shape species distributions, whilst minimizing the number of variables to allow modelling of species with few occurrence records. Variables included a number of bioclimatic parameters (1,2,4,5,11,12,13,14,15: http://worldclim.org/version2) in addition to productivity and other climate metrics (NDVI, seasonality, actual evapotranspiration, potential evapotranspiration seasonality and mean annual potential evapotranspiration, aridity, Emberger’s pluviotonic quotient, continentality, thermicity, maximum temperature of the coldest month - http://envirem.github.io/) - and both NDVI seasonality and mean). In addition, we included some topographic variables including soil pH, distance to bedrock, average tree height and tree density. All variables were clipped to a mask of Tropical Southeast Asia and southern China at a resolution of 0.008 decimal degrees (approximately 1km^2^) in ArcMap 10.3, then converted to asci format for modelling.

Models of Rhinolophid diversity were run in Maxent 3.4.4. Five replicates were run for each species, and the average taken before reclassifying with the 10^th^ percentile cumulative logistic threshold to form binary maps for each species (see Hughes et al., 2013). AUC for training and testing was 0.96 and 0.92 respectively, and all training AUCs were above 0.88.

Because of complex regional biogeography, optimal species habitat can exist in areas that have not been colonized. Therefore, we downloaded mapped ranges for 39 of the 49 species modelled from the IUCN (https://www.iucnredlist.org/resources/spatial-data-download). Bats were extracted from this data, clipped to match the study area. We then divided the IUCN data into five regions; mainland Southeast Asia, Philippines, Java-Sumatra, Borneo and Sulawesi-Moluccas, using shapefiles of each region to clip out bats listed there. This was collated to form a spreadsheet listing each zone each species was listed in, and then the appropriate shapefiles used to determine the ranges of each species (although only 39 of the 49 species could be treated in this way). Each species was then remosaiced with the mask to provide a binary distribution map, removing any potentially suitable areas that were outside the species biogeographic range. Stricter filters were not used, because for the majority of species there is not a clear analysis of genuine delineations of species ranges of if these species are migratory. These binary mosaicked maps were then summed with the other ten species using the mosaic tool to generate a map of richness for the region.

### QUANTIFICATION AND STATISTICAL ANALYSIS

No statistical analyses were conducted as part of this study.

## Notes

### Competing Interest Statement

The authors have declared no competing interest.

